# HypoAD: volumetric and single-cell analysis reveals changes in the human hypothalamus in aging and Alzheimer’s disease

**DOI:** 10.1101/2025.10.08.681270

**Authors:** Doudou Yu, Jürgen Germann, Ghulam Murtaza, Kaitlyn H. Hajdarovic, Kelsey R. Babcock, Shiva Kazempour Dehkordi, Alexander C. Jackson, Ivana Delalle, Miranda E. Orr, Habil Zare, Ritambhara Singh, William Stafford Noble, Andrei G. Vlassenko, Manu S. Goyal, Ashley E. Webb

## Abstract

Alterations in metabolism, stress response, sleep, circadian rhythms, and neuroendocrine processes are key features of aging and neurodegeneration. These fundamental processes are regulated by the hypothalamus, yet how its functionally distinct subregions and cell types change during human aging and Alzheimer’s Disease (AD) remains largely unexplored. Here, we present **HypoAD**, a comprehensive atlas of the human hypothalamus in aging and AD, integrating high-resolution MRI from 202 individuals with single-nucleus RNA-seq (snRNA-seq) of 614,403 nuclei from young, AD, and age-matched non-dementia controls. Our analysis reveals that hypothalamic subregions governing metabolism, stress, and circadian rhythms are particularly vulnerable, exhibiting significant changes in both volumes and gene expression during aging and AD. At the molecular level, machine learning models identified the inflammatory response and regulators of circadian rhythms as key cellular predictors of AD. These signatures were reflected in specific cell types: microglia transitioned to a pro-inflammatory state, while inhibitory neurons within sleep-and circadian-regulating hypothalamic subregions showed the most profound transcriptional alterations, including disruptions in ligand-receptor interactions and G-protein-coupled receptor signaling. Together, HypoAD provides a high-resolution volumetric map and a comprehensive transcriptomic atlas of the human hypothalamus in aging and AD, linking lifestyle and behavioral changes to their underlying volumetric and molecular pathways. Additionally, HypoAD provides a framework to investigate hypothalamic dysfunction and establishes a roadmap for targeted interventions aimed at mitigating physiological disruptions to potentially slow disease progression.

## Introduction

Alzheimer’s Disease (AD) is an age-associated neurodegenerative disease affecting millions worldwide, with a prevalence of approximately 10-30% among individuals over 65 years old^1^. Although its cardinal feature is cognitive impairment, AD manifests as a systemic condition involving profound disruptions in sleep, circadian rhythms, metabolic homeostasis, stress responses, and neuroendocrine functions^2^. The canonical neuropathological hallmarks are extracellular amyloid-β plaques and intracellular neurofibrillary tangles (NFT), which alongside widespread neuroinflammation and volumetric brain changes, also intensify with normal aging^2–7^. Because cognitive decline has traditionally guided both diagnosis and research, investigation has mainly focused on the hippocampus and cortex^8,9^. As the effects of AD on other brain regions remain poorly understood, investigating these understudied areas is crucial for elucidating the full spectrum of disease mechanisms and developing novel therapeutics^10,11^.

The hypothalamus, a small but complex brain region, orchestrates a wide range of physiological and neuroendocrine processes frequently dysregulated in aging and AD^12,13^. Emerging evidence suggests that hypothalamic dysfunction not only correlates with but also actively exacerbates AD pathology and cognitive decline^2^. Specifically, pathology within hypothalamic subregions, such as the suprachiasmatic nucleus (SCN)^14–17^, arcuate nucleus (ARC)^2,18^, and mammillary body (MN)^19^, has been associated with prominent sleep disturbances, metabolic dysfunction, and memory deficits observed in AD patients. These systemic disruptions, in turn, are hypothesized to create a vicious cycle that further accelerates disease progression and cognitive decline^2^. Despite its central role in homeostasis and brain aging^13,20^, the hypothalamus remains an understudied region in the context of human aging and AD. In particular, we have a limited understanding of the volumetric, cellular, and molecular changes within the human hypothalamus during these processes. While studies in animal models have highlighted hypothalamic involvement in aging and neurodegeneration^13,20–22^, human studies remain scarce. This paucity is especially pronounced for investigations that combine high-resolution imaging with single-cell analyses, complementary approaches essential for dissecting the anatomical and cellular heterogeneity of this complex brain region. This knowledge gap hinders our ability to determine how specific hypothalamic subregions and cell populations contribute to aging and AD, and why they exhibit differential vulnerability.

Here, we present HypoAD, a multi-modal atlas combining two powerful, cross-scale approaches. First, we performed high-resolution volumetric analysis of clinical imaging data from 202 individuals across the lifespan with varying cognitive and pathological statuses. Second, we generated a complementary comprehensive single-cell transcriptomic atlas, comprising 614,403 nuclei from the hypothalami of 17 post-mortem donors, including young and aged individuals with or without clinical and neuropathological diagnoses of AD. Integrating these datasets reveals the subregion-and cell-type-specific alterations of the human hypothalamus in aging and AD.

## Results

### Sex-specific alterations in hypothalamic subregion volumes with aging and AD

Neuroimaging studies have reported changes in hypothalamic volumes associated with aging and cognitive decline^23,24^. However, previous analyses were relatively low resolution and limited to a few major hypothalamic areas, thereby masking changes within structurally and functionally distinct subregions relevant to disease, such as the arcuate nucleus (AN) and suprachiasmatic nucleus (SC)^24^. To overcome this limitation, we leveraged a newly developed volumetric atlas of the human hypothalamus^25^ to delineate and measure 13 hypothalamic and 12 extrahypothalamic structures (Supplementary Table 1). Our analysis included a clinical imaging dataset of 202 individuals. The cohort had a mean age of 62.7 ± 14.2 years (mean ± standard deviation) and included 54.0% women, 10.4% with abnormal cognitive status, and 19.3% who were amyloid-β positive. The mean total hypothalamic volume for this cohort was 738.4 ± 76.4 mm^3^.

Pearson correlation analysis revealed that while the total hypothalamic volume showed no significant change with age (*r* = 0.031, adjusted *p* = 0.69), its constituent subregions displayed divergent age-related trajectories. This highlights that analyzing the hypothalamus as a whole can obscure distinct underlying patterns. Specifically, the lateral hypothalamus (LH) volume was negatively correlated with age (*r* = -0.41, adjusted *p* = 2.4e-9), suggesting age-related atrophy. In contrast, the volumes of the SC (*r* = 0.27, adjusted *p* = 0.00018), AN (*r* = 0.31, adjusted *p* = 1.1e-5), paraventricular nucleus (Pa) (*r* = 0.32, adjusted *p* = 4.6e-6), periventricular nucleus (Pe) (*r* = 0.40, adjusted *p* = 5.5e-9), and dorsomedial hypothalamus (DM) (*r* = 0.20, adjusted *p* = 0.0063) all increased with age with age across all 202 individuals (Fig. 1a, Supplementary Table 2). For comparison, most extrahypothalamic structures showed age-related volume reductions (Extended Data Fig. 1a, Supplementary Table 2). Notably, the positive correlation between DM volume and age was absent in cognitively normal, amyloid-β negative individuals (Extended Data Fig. 1b, Supplementary Table 2. This suggests that age-related changes in the DM, a subregion implicated in sleep-wakefulness^26^ and body weight regulation^27^, might be specifically linked to neurodegenerative processes rather than normal aging.

**Figure 1:**
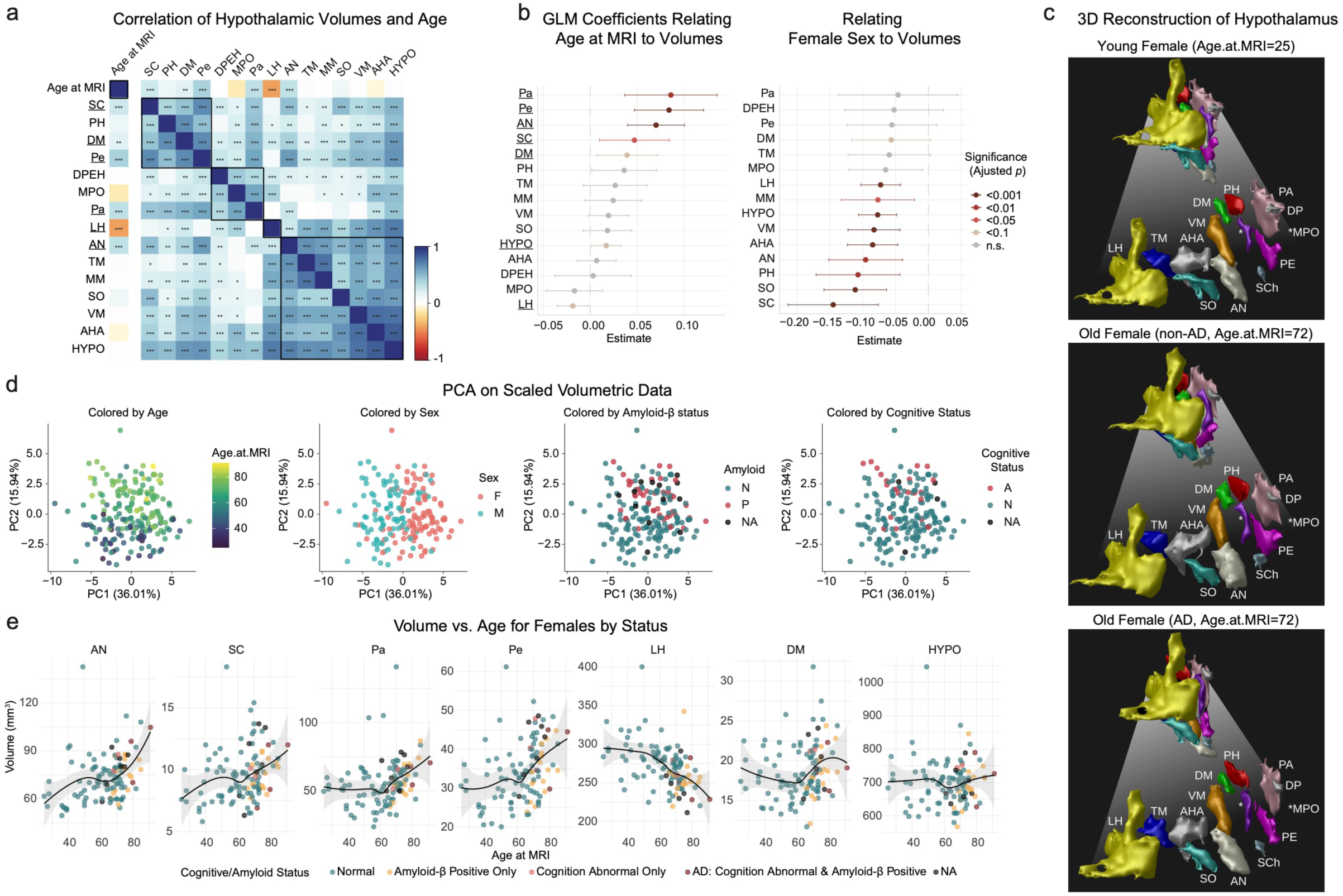
Sex-specific alterations in hypothalamic subregion volumes with aging and AD. **a,** Pearson correlation heatmap of hypothalamic subregion volumes with age at MRI. Blue and red indicate positive and negative correlations, respectively. Significance was determined after Benjamini-Hochberg (BH) correction (\**p* < 0.05, \*\**p* < 0.01, \*\*\**p* < 0.001). **b,** Forest plots showing GLM coefficients for the effects of age at MRI (left) and biological sex being female (right) on hypothalamic subregion volumes. Points are coefficient estimates and error bars represent 95% confidence intervals, with significance based on BH-corrected *p*-values. **C**, 3D reconstructions of hypothalamic subregions from MRI scans of three representative female individuals. **d,** PCA of hypothalamic subregion volumes. Points represent individual participants, positioned by the first two principal components (PC1 and PC2). The plots show participants colored by key variables age at MRI, sex, amyloid-β status (N, Normal; P, Positive), and cognitive status (N, Normal; A, Abnormal) **e,** Scatterplots depicting age-related volumetric changes (mm^3^) in selected hypothalamic subregions and the total hypothalamus in females. Each point represents an individual and is colored by their combined cognitive and amyloid-β status. A locally estimated smoothing (LOESS) curve (black line) and its 95% confidence interval (shaded area) are shown for each plot.

To account for covariates while evaluating the relationship between age and subregion volumes, we applied a generalized linear model (GLM). After controlling for biological sex, intracranial volume (ICV), cognitive status, and amyloid-β pathology, LH volume showed a negative association with age (adjusted *p* = 0.069). In contrast, the SC (adjusted p = 0.047), AN (adjusted *p* = 5.6e-5), Pa (adjusted *p* = 0.0039), Pe (adjusted *p* = 7.2e-5), DM (adjusted *p* = 0.062), and total hypothalamic (HYPO) volumes (adjusted *p* = 0.084) showed a positive association with age (Fig. 1b, Extended Data Fig. 1c, Supplementary Table 3). Biological sex was also a significant factor; females exhibited smaller volumes in several subregions, most prominently in the SC (adjusted *p* = 0.00030, Fig. 1b, Extended Data Fig. 1c, Supplementary Table 3). This finding aligns with previous studies emphasizing the sex-dimorphic nature of the hypothalamus, including the SC^13,24,28^. Furthermore, abnormal cognitive status was associated with larger volumes in the DM (adjusted *p* = 0.025) and Pe (adjusted *p* = 0.068, Extended Data Fig. 1d, Supplementary Table 3), suggesting a link between volumetric changes in these regions and cognitive decline. As expected, most subregion volumes were positively correlated with ICV (Extended Data Fig. 1e, Supplementary Table 3). To visualize changes with age and AD status, hypothalamic subregions from MRI scans of three representative female individuals were reconstructed in 3D (Fig. 1c).

To holistically evaluate volumetric relationships, we performed principal component analysis (PCA) on volumetric data from all 25 delineated hypothalamic and extrahypothalamic structures. This analysis revealed that the first two principal components (PCs) were primarily driven by age, sex, amyloid-β status, and cognitive status, causing individuals to cluster along these axes (Fig. 1d). Examination of volumetric trajectories via scatterplots stratified by cognitive and amyloid-β status revealed subregion-specific patterns that also differed by sex (Fig. 1e, Extended Data Fig. 1f). Notably, individuals with AD (defined as cognitive decline with Clinical Dementia Rating^29^ greater than 0 and amyloid-β positivity) displayed reduced LH volumes compared to their age-matched non-AD counterparts, consistent with its overall age-related decline (Extended Data Fig. 1g-h). In contrast, Pa, DM, and total HYPO volumes exhibited a further decline in individuals with AD compared to age-matched non-AD individuals, a pattern particularly evident in females (Extended Data Fig. 1h). Although paired Wilcoxon signed-rank tests were not statistically significant after correction, we observed a consistent pattern of small to moderate effect sizes for AD-related volume differences across subregions (AN=0.008, DM=0.277, Pa=0.176, Pe=0.134, SC=0.050, HYPO=0.209; LH=0.461).

These subregion-specific volumetric analyses provide high-resolution insights into hypothalamic aging trajectories and sex-dependent alterations, underscoring the importance of examining individual subregions. Our data confirm that aging is associated with volumetric changes in distinct subregions, even after controlling for amyloid-β deposition and cognitive status. The volumetric changes linked to cognitive impairment were particularly evident in the DM. Given that the DM receives major input from the SC^26^, the brain’s central pacemaker, this finding suggests that hypothalamic subregions governing circadian rhythms may be especially vulnerable to aging and age-related neurodegeneration.

### Single-cell analysis of the AD hypothalamus

While volumetric data provides anatomical context, it cannot reveal the cell-type-specific molecular changes that offer cell-type-specific insights into the aging and AD hypothalamus. To investigate such changes, we performed snRNA-seq on postmortem hemi-hypothalamus tissue from 14 individuals (see cohort details in Fig. 2a-b and Extended Data Fig. 2a). The cohort consisted of seven individuals with mostly early-stage AD (n=6 sporadic, n=1 familial) and age-matched, cognitively and neuropathologically normal controls (CTL) (n=7).

**Figure 2:**
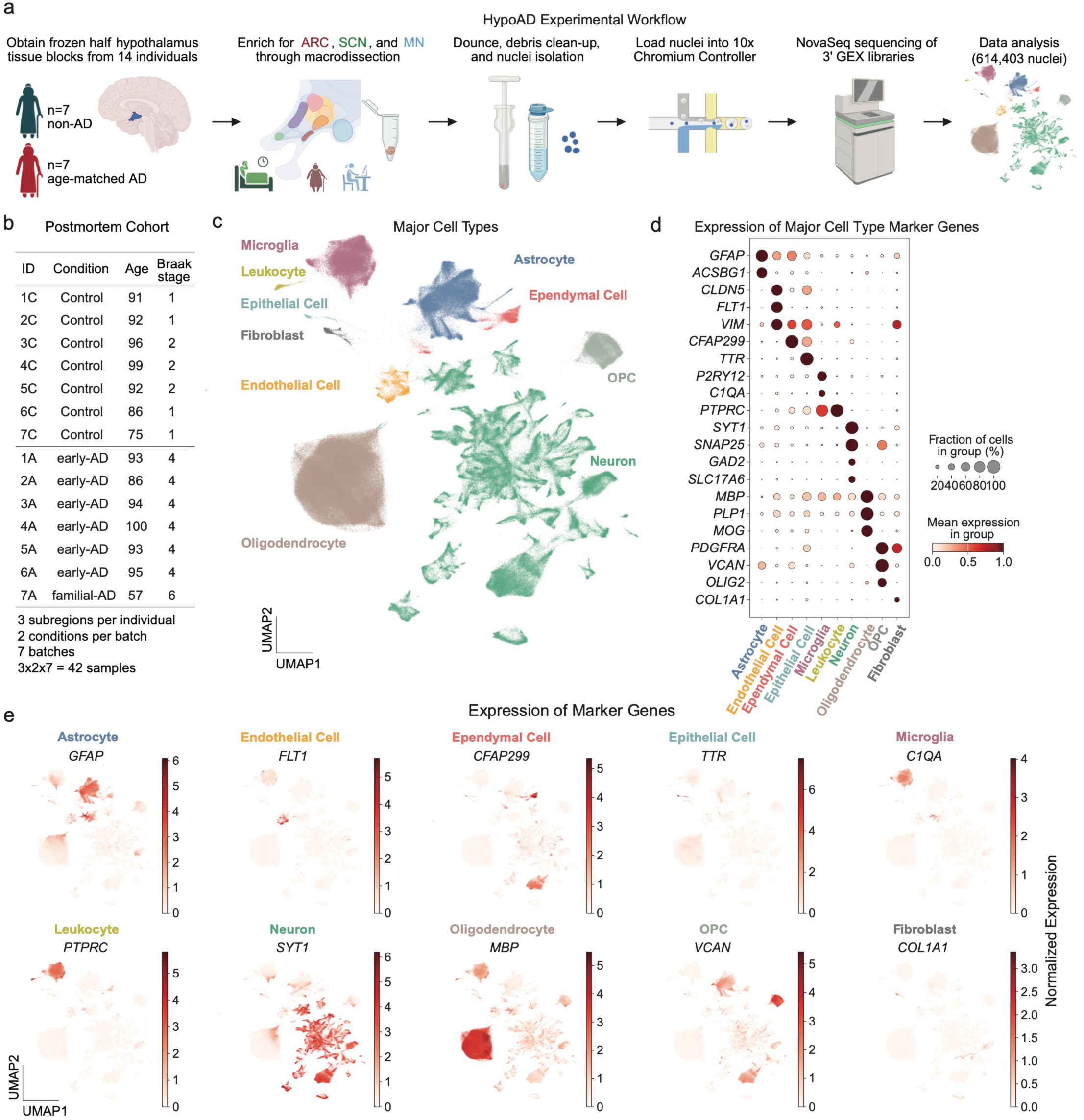
Single-cell analysis of the AD hypothalamus. **a,** Schematic of the single-nucleus RNA-sequencing (snRNA-seq) workflow. Frozen post-mortem hypothalamic tissue from 14 individuals (7 non-AD controls, 7 age-matched AD cases) was macrodissected to enrich for the ARC, SCN, and MN subregions. Nuclei were isolated, and libraries were prepared and sequenced on a NovaSeq platform. The resulting data were integrated with 133,879 publicly available nuclei from three young, healthy controls, yielding a final dataset of 614,403 nuclei across young control, aged control (non-AD), and AD groups for comparative analysis. **b,** Overview of the 14 individuals included in the snRNA-seq study, detailing their sample IDs, clinical condition (non-AD control, early-stage AD or familial AD), age at death, and Braak stage. Each individual contributed tissue from three hypothalamic subregions (ARC, SCN, MN), totaling 42 tissue samples processed across 7 experimental batches. **c,** UMAP plot of the integrated 614,403 nuclei. Distinct clusters represent identified major cell types, including neurons, astrocytes, oligodendrocytes, microglia, endothelial cells, ependymal cells, fibroblasts, epithelial cells, OPCs, and leukocytes. Each cell type is annotated and highlighted with a unique color. **d,** Dot plot of selected marker genes. The size of each dot corresponds to the fraction of cells within a given cell type expressing the marker, while the color intensity indicates the average normalized expression level. **e,** Normalized expression of key canonical marker genes mapped onto the UMAP plot from panel (**c**). Representative markers include *GFAP* (astrocytes), *FLT1* (endothelial cells), *CFAP299* (ependymal cells), *TTR* (epithelial cells), *C1QA* (microglia), *PTPRC* (CD45, leukocytes), *SYT1* (neurons), *MBP* (oligodendrocytes), *VCAN* (OPCs), and *COL1A1* (fibroblasts).

CTL individuals had no history of cognitive deficits or neurological diseases. The AD individuals predominantly presented with sporadic, clinically confirmed cognitive decline and were characterized by early-stage pathology (Braak stage IV; Fig. 2b) at autopsy. This stage is considered optimal for detecting significant transcriptional changes while sufficient neuronal populations are still present for analysis^30^. All samples exhibited high quality, with short postmortem intervals (PMIs) of less than nine hours and RNA integrity number (RIN) scores above 6.5. Importantly, hierarchical clustering showed that samples grouped by pathology status rather than by potential confounding factors such as time of death or technical quality metrics (Extended Data Fig. 2a).

Macrodissection was performed on the tissue blocks to enrich for the arcuate nucleus (ARC), suprachiasmatic nucleus (SCN), and mammillary nucleus (MN). Single nuclei were subsequently isolated in a cold room environment, and 3’ gene expression libraries were constructed and sequenced. Following rigorous quality control, a total of 480,524 nuclei were retained for downstream analysis, with an average of 9,639 ± 19.6 (mean ± Standard Error of the Mean, SEM) unique molecular identifier (UMI) counts and 3,000 ± 3 (mean ± SEM) detected genes per nucleus.

To enhance cell type annotation, we integrated our dataset with 133,879 previously published, high-quality human hypothalamic nuclei from three younger individuals (aged 29, 42, and 50 years)^31,32^ using the scVI algorithm^33^. The resulting integrated atlas is visualized as a Uniform Manifold Approximation and Projection (UMAP) plot (Fig. 2c), where cells are colored by major cell types identified through the expression of canonical marker genes (Fig. 2d-e; see Extended Data Fig. 2b-g for data quality control).

### Machine learning nominates inflammatory and circadian genes as key features of AD in the hypothalamus

To identify transcriptomic features that distinguish hypothalamic cells from individuals with AD versus controls, we trained and tested elastic net models on expression of 36,600 genes from 480,542 cells. The dataset was collected across seven independent batches, each containing one AD and one control subject (Fig. 3a). To ensure robustness and prevent data leakage, we implemented a leave-one-batch-out nested cross-validation (CV) scheme. For each of the seven outer folds, an inner 6-fold CV was used to tune the model’s hyperparameters. This approach generated seven distinct models, from which we ranked gene-associated weights to identify the most predictive features (Fig. 3a).

**Figure 3:**
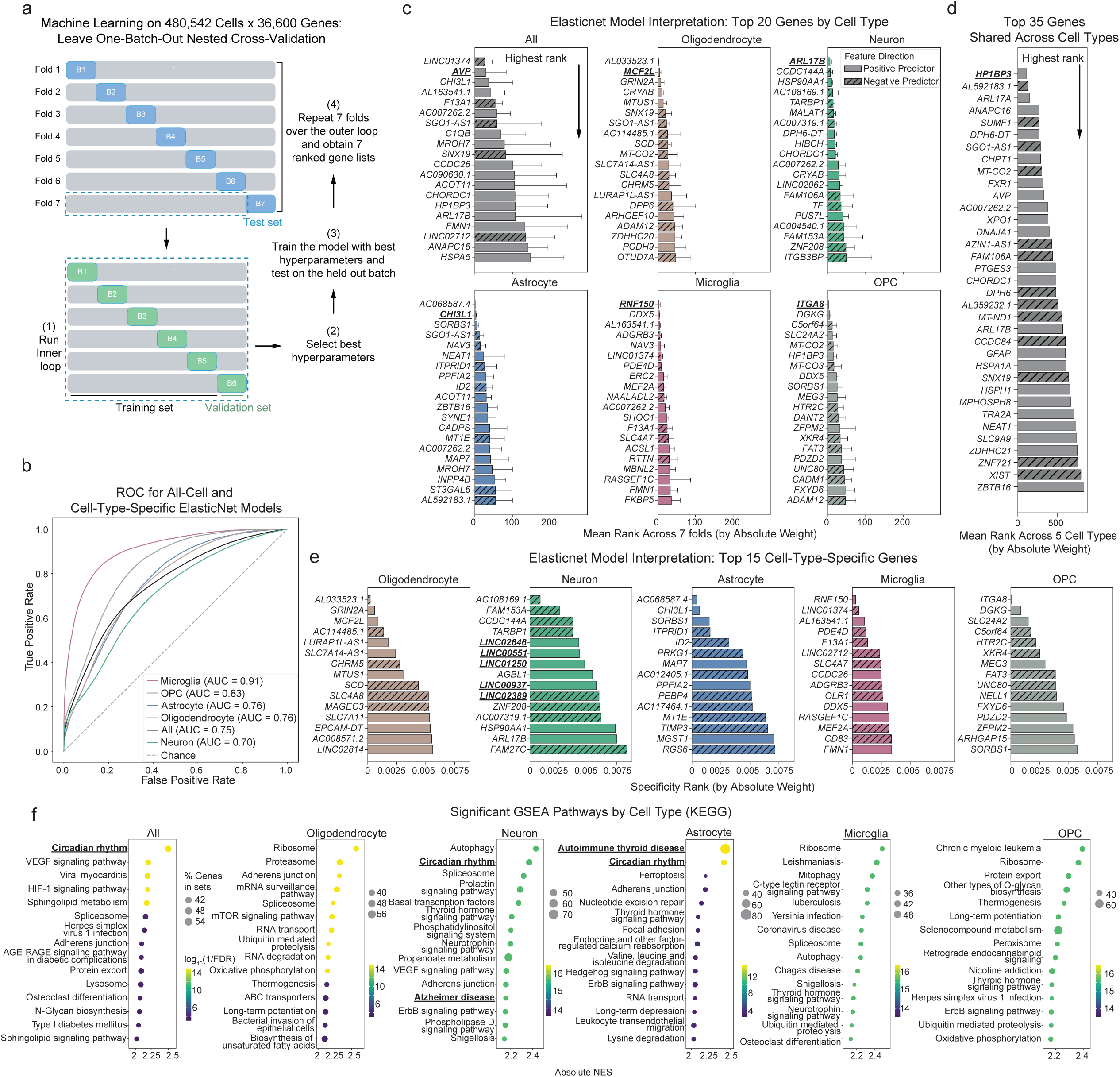
Machine learning models identified shared and cell-type-specific predictive features of AD. **a,** Schematic of the nested cross-validation strategy. An outer loop using leave-one-batch-out cross-validation isolated a single batch for testing, while an inner loop, also using leave-one-batch-out, selected optimal hyperparameters from the remaining training batches. **b,** ROC curves for the Elastic Net model trained on all cells combined (all-cell model) and on each of the five most abundant cell types. The AUC represents the pooled performance across all seven outer cross-validation folds. **c,** Top 20 predictive features (genes) for AD classification, ranked by the mean absolute weight from the Elastic Net model trained on all cells and on individual cell types. Bar plots show the mean rank across seven cross-validation folds; error bars represent the standard deviation. Solid bars indicate a positive predictive weight, while hatched bars indicate a negative predictive weight. A lower Specificity Rank value indicates higher feature importance. **d,** Top 35 shared features identified by their average rank across all five cell-type-specific models. The bar plot indicates the mean rank, with lower values representing higher importance. **e,** Top 15 cell-type-specific features for each cell type. Specificity was determined by the gene’s rank in its target cell type compared to its rank in all other cell types. Bar plot shows the specificity score. **f,** Selected significantly enriched KEGG pathways from GSEA. The analysis was performed on genes ranked by their signed model weights. The dot plot shows the top pathways for the all-cell model and for each of the five cell-type-specific models. The x-axis represents the absolute normalized enrichment score (NES), dot size represents the number of genes in a pathway, and color indicates to the significance (−log10(FDR)).

We first trained a model on all cells, which achieved predictive performance (average AUROC = 0.70). We then trained models specific to the five most abundant cell types. All cell-type-specific models also performed well, with the microglia model showing the highest accuracy (average AUROC = 0.91). The comparatively lower performance for neuronal models may be attributable to the complexity and heterogeneity of neurons in the hypothalamus (Fig. 3b).

To interpret our models, we averaged the absolute gene weights from the seven outer CV folds to identify the top 20 most predictive features for the all-cell and cell-type-specific classifiers (Fig. 3c). Notably, many top-ranked genes have been previously implicated in AD pathology, lending biological support to our model’s predictions. For instance, the all-cell model’s highest-ranked features included arginine–vasopressin (AVP), a hypothalamic hormone and neurotransmitter involved in circadian rhythms whose concentration is reportedly elevated in AD patients^34–36^. For our cell-type-specific models, top genes also had clear links to AD. The oligodendrocyte model highlighted *MCF2L*, a gene with known differential methylation in AD^37^ that was also a significant hit in epigenome-wide association studies^38,39^. The neuron model identified ARF Like GTPase 17B (*ARL17B*), a cell-type-specific expression quantitative trait locus (eQTL) that shows elevated expression in the prefrontal cortex of AD patients, particularly in inhibitory neurons^40^. In astrocytes, a key feature was Chitinase-3-like protein 1 (*CHI3L1*), an astrocyte-secreted protein that mediates neuroinflammation and serves as a cerebrospinal fluid biomarker for early AD diagnosis^41^. In microglia, Ring Finger Protein 150 (*RNF150*) was identified as a significant cis-eQTL^42^. Finally, the OPC model prioritized Integrin Subunit Alpha 8 (*ITGA8*), a gene linked to demyelination^43^, which is a pathological feature of AD (Fig. 3c).

We next sought to distinguish predictive genes that were shared across cell types from those that were cell-type-specific. By calculating the mean rank of genes across the five most abundant cell types, we identified heterochromatin protein 1 binding protein 3 (*HP1BP3*) as the top shared feature (Fig. 3d). *HP1BP3* is a candidate modulator of cognitive aging in the hippocampus, and its levels are affected by metformin, a drug under investigation in anti-aging clinical trials^44,45^. However, its function in the hypothalamus remains unexplored. To identify cell-type-specific features, we calculated a specificity rank for each gene (Fig. 3e). This approach uniquely revealed a strong enrichment of long noncoding RNAs (e.g., *LINC02646*, *LINC00551*, *LINC01250, LINC00937,* and *LINC02389*) among the top-ranked genes for neurons, suggesting their potential role in AD (Fig. 3e). Together, this analysis identifies both previously documented and novel AD-associated genes, providing a rich, cell-type-specific resource to guide future hypothesis generation.

To understand the broader biological pathways implicated by our models, we performed a Gene Set Enrichment Analysis (GSEA) on the ranked predictive genes. This revealed a significant enrichment for gene sets involved in the regulation of circadian rhythms and inflammatory pathways (Fig. 3f, Extended Data Fig. 3, Supplementary Table 4), connecting our cell-level classifiers to established hallmarks of neurodegeneration.

### Cell-type-specific transcriptional changes

To complement our machine learning analysis, we performed cell-type-specific differential gene expression analysis by comparing pseudobulk profiles (aggregated by cell type and sample) from early-AD and control samples. This analysis, conducted with the edgeR likelihood ratio test (LRT) model, identified distinct transcriptional signatures for each major cell type (Fig. 4a, Supplementary Table 5). Notably, microglia exhibited the largest number of differentially expressed genes (DEGs), highlighting their substantial transcriptional response even in early AD.

**Figure 4:**
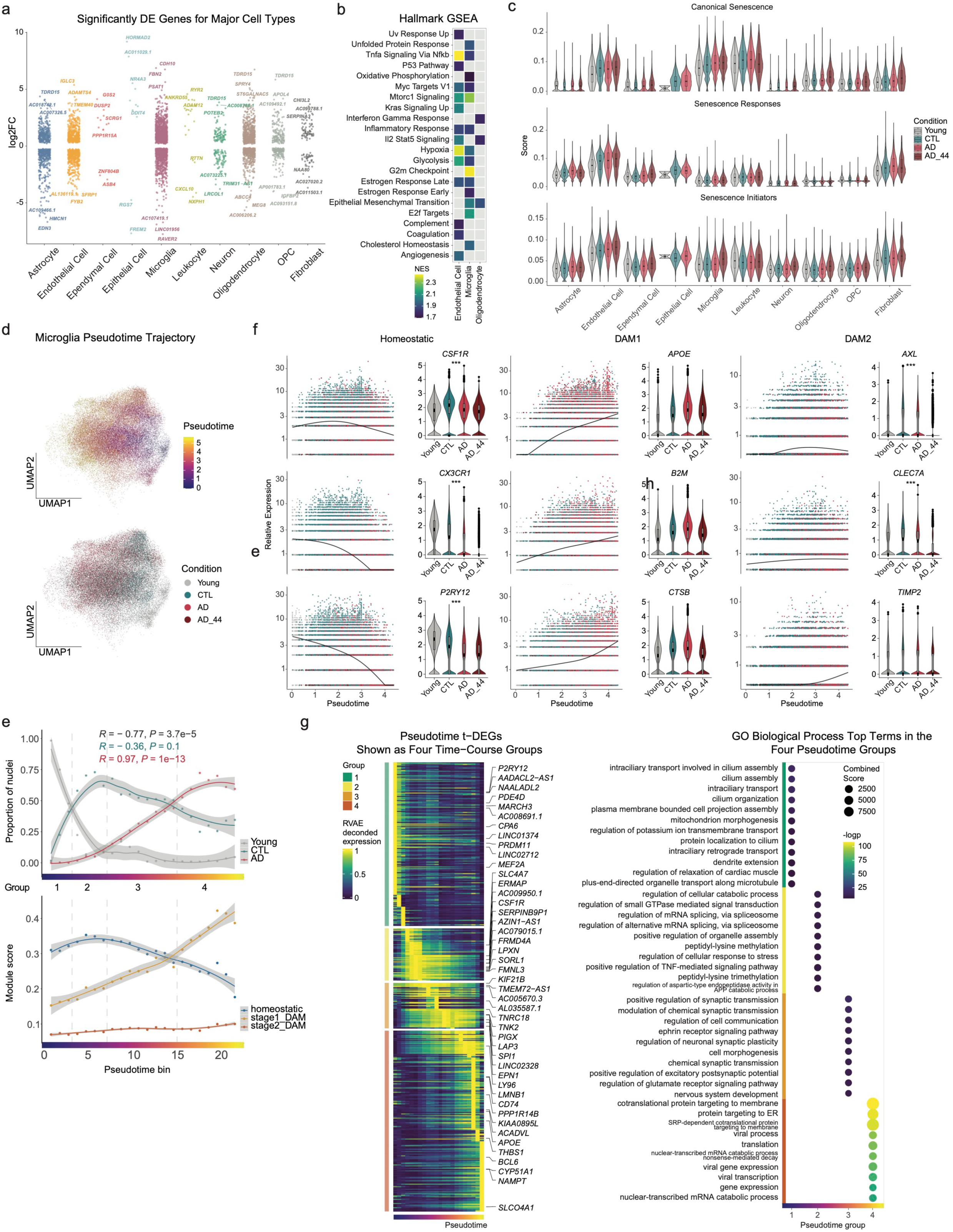
Cell-type-specific transcriptional changes and the microglial aging and disease trajectory in HypoAD. **a,** Strip plot illustrating significant DEGs (|log2FoldChange|>0 and FDR<0.05, identified using pseudobulk-edgeR-LRT) in various cell types when comparing early-AD with CTL samples. DEGs were identified using a pseudobulk approach with the edgeR likelihood-ratio test. **b,** Heatmap displaying significantly enriched Hallmark pathways in non-neuronal cell types associated with early-AD. **c,** Senescence scores across major cell types. Violin plots showing scores for three senescence-related gene signatures for major cell types. **d,** UMAP plots of microglial nuclei. The top UMAP is colored by inferred pseudotime, illustrating the continuous trajectory. The bottom UMAP is colored by condition: young, CTL, AD, and AD_44. **e,** Top: Scatterplots showing the proportion of microglial nuclei from each condition (young, CTL, AD, AD_44) across pseudotime. The trajectory is divided into four illustrative stages for visualization: aging-young (group 1), aging-aged CTL (group 2), disease-CTL (group 3), and disease-AD (group 4). Bottom: Line plot depicting the scores of homeostatic, DAM1, and DAM2 microglial signatures across the pseudotime trajectory, showing a progression from homeostatic states towards DAM1 and subsequently DAM2. **f,** Heatmap showing distinct clusters of t-DEGs identified by Moran’s *I* test, that exhibit coordinated temporal expression patterns across the microglial aging and disease trajectory. **g,** Dot plot showing significantly enriched GO BP for the distinct clusters of co-expressed t-DEGs from panel (f), highlighting key functional transitions along the trajectory.

To understand the functional implications of this response, we performed a GSEA on the microglial DEGs. This revealed a significant enrichment for pathways involved in G2M checkpoint regulation, mTORC1 signaling, and cholesterol homeostasis. These findings suggest that in early AD, microglia enter a complex, supportive state. The enrichment of mTORC1 signaling points to metabolic reprogramming and cellular activation, while the cholesterol homeostasis signature is consistent with enhanced cholesterol efflux—a process linked to amyloid-β phagocytosis. We propose that these coordinated processes reflect a neuroprotective microglial response to support neurons in the early-AD microenvironment (Fig. 4b).

Cellular senescence can contribute to aging and neurodegeneration by promoting chronic inflammation and disrupting tissue homeostasis^46–49^. We explored this by evaluating three established curated senescence signatures — Canonical Senescence pathways, Senescence Responses, and Senescence Initiators(Fig. 4c)—alongside three NFT signatures (Extended Data Fig 4. a-c), which have shown overlap with senescence-associated pathways^47^, across all major cell types. Microglia, leukocytes, fibroblasts, and endothelial cells exhibited the highest senescence and NFT signature scores, which were elevated with both aging (CTL group) and AD (Fig. 4c, Extended Data Fig 4. a-c). In contrast, neurons showed NFT enrichment that became particularly pronounced in AD individuals carrying the *APOE*4/4 genotype (denoted as AD_44) (Fig. 4c, Extended Data Fig 4.a-c). These findings suggest a dichotomy: senescence in non-neuronal cells may drive microenvironmental dysfunction in AD, while neuronal NFTs reflect a distinct, tau-driven pathology that escalates with disease progression or genetic susceptibility. This highlights the potential for different, cell-type-specific therapeutic targets.

### Microglia trajectory in HypoAD

Microglia have been extensively studied in brain aging and neurodegenerative diseases^20,50,51^. To investigate the extent to which similar changes occur in the aging and AD hypothalamus, we applied trajectory inference^52,53^ to map the continuum of microglial states. By integrating microglia from all conditions (young, aged CTL, and AD), we inferred a continuous trajectory representing a transition from aging to AD using Monocle3^52^ (Fig. 4d, Extended Data Fig. 4d). This pseudotime continuum was categorized into four main stages: an initial “aging phase” predominantly composed of microglia from young (stage 1) and aged CTL (stage 2) individuals, followed by a “disease phase” where microglia from aged CTL (stage 3, representing a later state for these cells along the trajectory) and AD (stage 4) individuals became more prevalent (Fig. 4e). Interestingly, a subset of microglia from aged CTL individuals appeared at earlier trajectory points, while some microglia from young individuals persisted into later stages, suggesting that certain microglia may retain youthful properties while others exhibit heightened susceptibility to early pathological changes.

To characterize microglial activation states along this inferred trajectory, we examined three established microglial gene signatures^54^: (1) homeostatic microglia, (2) stage 1 disease-associated microglia (DAM1), typically activated by aging and amyloid plaques, and (3) stage 2 DAM (DAM2), which is dependent on TREM2 signaling^54,55^. Using UCell^56^ to score these signatures across pseudotime, we observed a clear transition from a homeostatic state towards DAM1, accompanied by a modest increase in DAM2 signatures (Fig. 4e). Kinetic and violin plots of representative genes further supported this, demonstrating a gradual decline in homeostatic markers (e.g., *CSF1R*, *CX3CR1*), a marked increase in DAM1-associated genes (e.g., *APOE*, *B2M*), and a modest rise in DAM2 markers (e.g., *AXL*, *CLEC7A*) (Fig. 4f). This observed shift from homeostatic to DAM states underscores the critical role of microglial functional changes in AD progression and highlights the potential for therapeutic interventions aimed at preserving homeostatic functions or preventing the full transition to detrimental DAM states.

To capture the full spectrum of transcriptional changes along the inferred microglial pseudotime, we performed Moran’s *I* test, identifying 13,104 trajectory-associated differentially expressed genes (t-DEGs) (Supplementary Table 6, Extended Data Fig. 4e). To visualize their dynamic expression patterns, we applied RVAgene^57^, an autoencoder-based neural network framework, to reconstruct and smooth gene expression trends, which were then displayed as a heatmap (Fig. 4g). Gene Ontology (GO) biological process (BP) analysis of these t-DEGs revealed distinct transcriptional transitions: during the aging portion of the trajectory, BPs shifted from cilium assembly and mitochondrial morphogenesis towards stress responses, including amyloid precursor protein (APP) catabolic processes. Advancing into the AD portion, the transition progressed from cell morphogenesis and synaptic transmission modulation to viral response pathways and protein targeting to the endoplasmic reticulum (ER) (Fig. 4g).

### Neuronal diversity in HypoAD across subregions

Neurons in the human hypothalamus exhibit extensive cell type diversity^31,32,58^. We analyzed a total of 213,469 neuronal nuclei, classifying them into three major neurotransmitter-defined categories: inhibitory/GABAergic (expressing *GAD2*), excitatory/glutamatergic (expressing *SLC17A6*), and a smaller population of histaminergic neurons (expressing *HDC*) (Fig. 5a-c).

**Figure 5:**
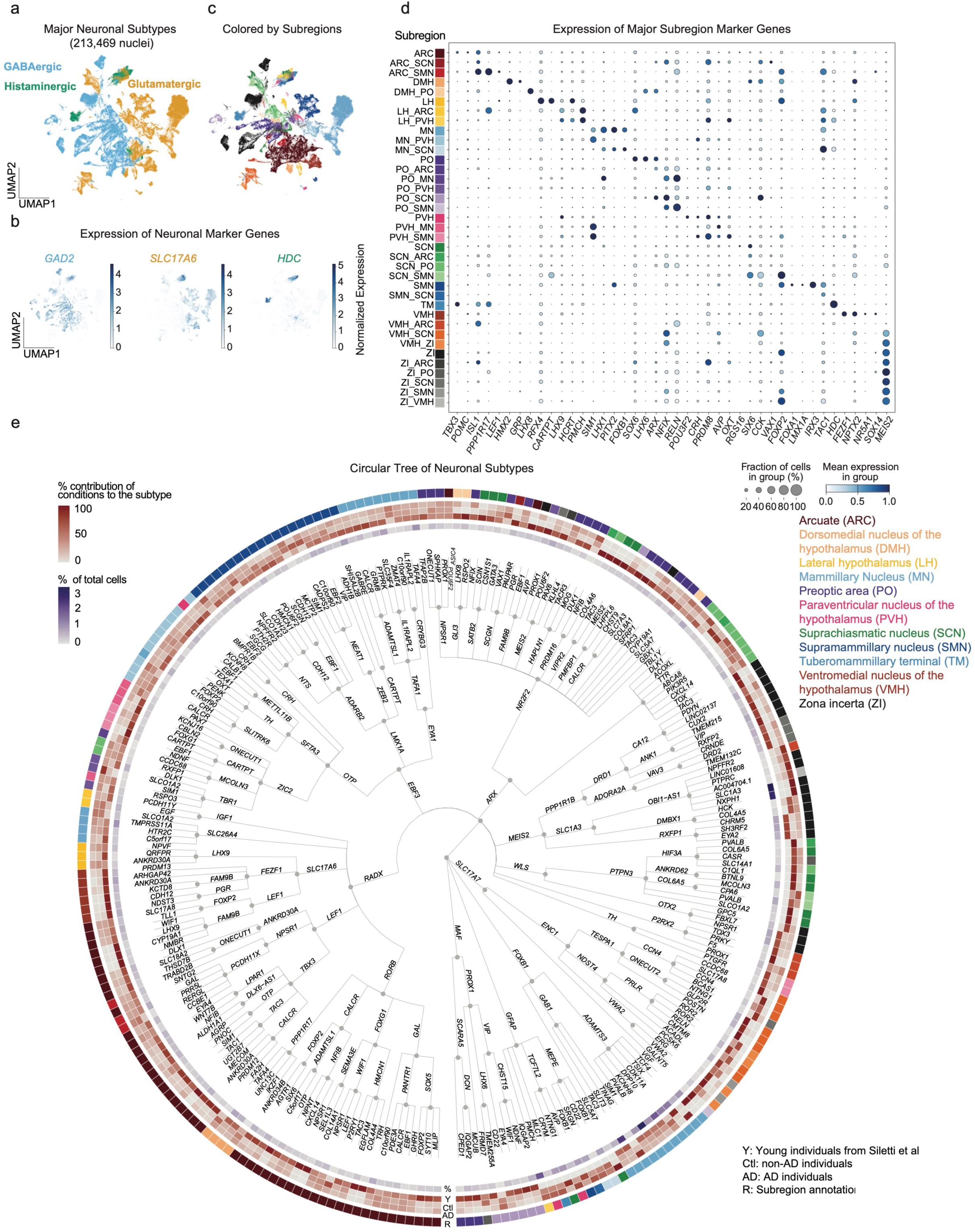
Neuronal diversity in HypoAD across hypothalamic subregions. **a,** UMAP of 213,469 hypothalamic neurons, color-coded by major neurotransmitter category: GABAergic (*GAD2*-expressing; blue), glutamatergic (*SLC17A6*-expressing; orange), and histaminergic (*HDC*-expressing; green). **b,** Feature plots overlaid on the UMAP, illustrating the normalized expression of *GAD2* (GABAergic), *SLC17A6* (glutamatergic), and *HDC* (histaminergic) across the neuronal populations. **c,** UMAP plot of neuronal nuclei, with cells colored according to their inferred subregional identity based on marker gene expression. This panel highlights the regional distribution of neuronal clusters within the hypothalamus. **d,** Dot plot displaying the expression of key marker genes enriched in distinct hypothalamic subregions as inferred by marker expression. The size of each dot represents the fraction of nuclei within a neuronal subtype expressing the gene, and the color intensity indicates the mean normalized (or scaled) expression level. **e,** A circular dendrogram illustrating the relationships among the 254 identified hypothalamic neuron subtypes. The dendrogram is annotated to show, for each subtype, its inferred subregional identity (based on marker genes), and its relative contribution from young, aged CTL and AD individuals.

To further delineate neuronal subtypes, Leiden clustering was applied at multiple resolutions. This hierarchical approach yielded five distinct levels of clustering, with the number of identified subclusters ranging from 5 at the broadest level to 254 at the finest resolution. We inferred the anatomical origin of these neuronal subclusters by leveraging a curated set of subregional marker genes^31^(Fig. 5d). Subclusters exhibiting markers for multiple subregions were annotated accordingly, with the primary subregion listed first based on dominant marker expression (Fig. 5d).

A hierarchical dendrogram was constructed to visualize the relationships among these neuronal subtypes across the different clustering resolutions. Nodes and leaves within this dendrogram were labeled with specific marker genes that distinguish each subtype from other subtypes at the same hierarchical level. The final set of 254 refined leaf nodes (subclusters) was comprehensively annotated with details including their relative abundance within the total neuronal population, their distribution across young, aged CTL, AD conditions, and their inferred subregional identity (Fig. 5e). We mapped the 254 neuronal subtypes to Visium spatial transcriptomics data from nine human donors to validate their inferred cell type identities (Extended Data Fig. 5)^58,59^. This detailed characterization provides a robust foundation to understanding how specific neuronal subtypes are altered under different physiological and pathological conditions.

### Neuronal subtype-specific transcriptional signatures in HypoAD

The hypothalamus comprises distinct neuronal populations with specialized homeostatic roles, yet their subtype-specific contributions to AD remain poorly characterized. To address this, we performed differential gene expression analysis comparing early-AD to CTL groups using the edgeR LRT model on pseudobulk expression profiles aggregated by neuronal subtype and sample. This identified early-AD-associated transcriptional signatures across 254 distinct human hypothalamic neuronal subtypes. We prioritized the top 30 subtypes exhibiting the most significant numbers of DEGs; these DEGs were then grouped by their classification into major neuronal categories (GABAergic, glutamatergic, or histaminergic) for visualization (Fig. 6a). Notably, GABAergic/inhibitory neurons displayed the most pronounced transcriptional changes. Enriched Reactome pathways within these neurons highlighted alterations in peptide-ligand binding receptors and G-protein coupled receptor (GPCR) signaling (Fig. 6b, Supplementary Table 7), suggesting a potential dysregulation of neuropeptide signaling and synaptic function in the early-AD hypothalamus that may contribute to metabolic disturbances characteristic of AD pathogenesis.

**Figure 6:**
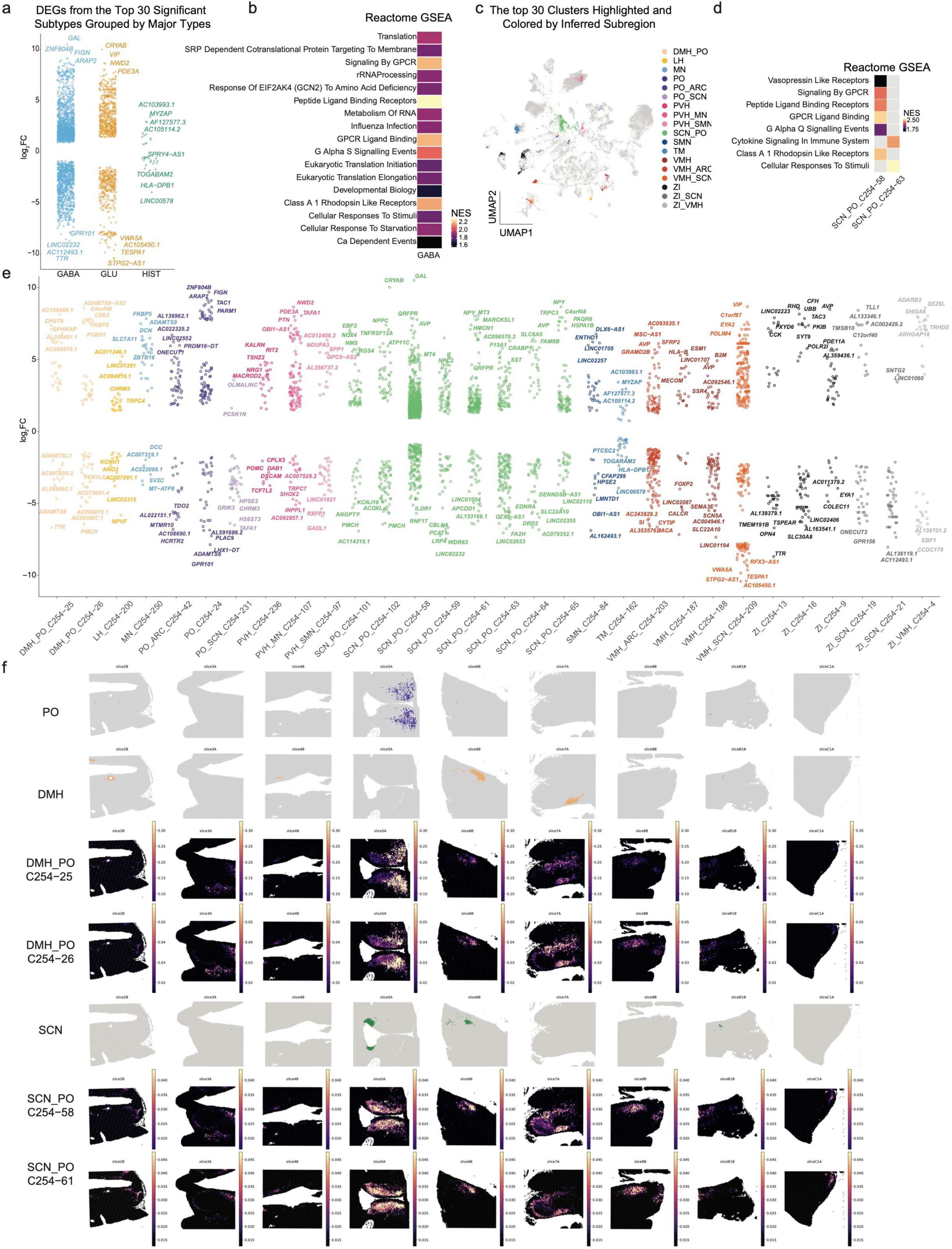
Neuronal subtype-specific transcriptional signatures in HypoAD. **a**, Significant DEGs identified within the top 30 transcriptionally altered neuronal subtypes. DEGs are grouped by whether the subtype belongs to the GABAergic, glutamatergic (GLU), or histaminergic (HIST) neuronal category. **b,** Heatmap displaying significantly enriched Reactome pathways within AD-altered GABAergic neurons **c,** UMAP plot showing the 30 neuronal subtypes with the most significant transcriptional changes in early AD. Subtypes are colored by their inferred hypothalamic subregional origin. **d,** Heatmap displaying significantly enriched Reactome pathways within AD-altered neuronal subtypes. Highlighted pathways include those related to vasopressin-like receptors, GPCR signaling, and immune/cytokine signaling. **e,** Strip plot showing log2FC values for significant DEGs identified across all 254 neuronal subtypes. Points or strips can be colored to indicate the predominant inferred hypothalamic subregion of the neuronal subtype in which the DEG is found. **f,** Inferred spatial localization of top AD-altered neuronal subtypes. Visium spots are colored by the predicted proportion of each indicated subtype, mapping transcriptional changes to anatomical space.

The top 30 transcriptionally vulnerable neuronal subtypes were further categorized by their inferred subregional origins and visualized in UMAP embeddings and spatial locations (Fig. 6c-f, Extended Data Fig. 6a). This analysis highlighted neurons within the SCN as particularly vulnerable, exhibiting substantial transcriptional alterations in AD.

For instance, Reactome pathway analysis of a prominent SCN neuronal cluster (C254-58) revealed significant changes in GPCR signaling and vasopressin-related receptor pathways. Vasopressin, one of the most abundant neuropeptides in our dataset (Extended Data Fig. 6b), was also identified as the top feature in our ML analysis. This key neuropeptide is primarily synthesized and secreted by SCN neurons in a circadian manner^60^ and is crucial for maintaining circadian homeostasis. These findings suggest that early-AD disrupts SCN neuronal function, potentially contributing to the circadian dysregulation often observed in patients.

Together, these analyses reveal selective transcriptional dysregulation in specific hypothalamic neuronal subtypes and subregions in AD, with a particular impact on GABAergic/inhibitory neurons and the SCN. The pronounced alterations in neuropeptide signaling (e.g., vasopressin) and GPCR pathways within the SCN suggest a mechanistic link to circadian rhythms disruption, a hallmark of AD that can precede cognitive decline. These findings position the hypothalamus as a critical, vulnerable hub in AD pathogenesis, advocating for therapeutic strategies that target neuropeptide-GPCR signaling, circadian entrainment, or subregion-specific resilience mechanisms to potentially mitigate preclinical AD symptoms.

## Discussion

Our findings provide a comprehensive, multimodal view of how the hypothalamus is altered in aging and early AD, illustrating region- and cell-type-specific changes that have not been observed previously. Neuroimaging studies have suggested that hypothalamic volume correlates with both aging and cognitive decline. Here, by utilizing the highest-resolution atlas available of the human hypothalamus^25^, we detected specific volumetric changes in thirteen distinct hypothalamic subregions that would be masked by more coarse-grained analyses. Our results show that while certain nuclei (e.g., the lateral hypothalamus, LH) diminish in size with age, others (e.g., suprachiasmatic nucleus (SCN), arcuate nucleus (AN), paraventricular nucleus (Pa), periventricular nucleus (Pe), and dorsomedial hypothalamus (DM)) can exhibit an increase in volume among older adults, even when controlling for amyloid-β pathology and cognitive impairment. Moreover, PCA of volumetric data underscored substantial sex differences, consistent with prior evidence that hypothalamic structure and function differ markedly between males and females^24,28^. Cognitive impairment was further associated with differences in DM, linking dysregulation of circadian rhythms to neurodegeneration. However, in comparison to aging and sex, differences in the volumes of hypothalamic nuclei in the context of amyloid-β deposition and cognitive impairment were relatively modest, which could be related to lower AD-related neurodegeneration in the hypothalamus compared to other brain regions or, instead, a poor sensitivity of volumetric analysis in detecting AD-related effects upon the hypothalamus with limited sample size.

To investigate the cellular and molecular effects of AD in individual hypothalamic nuclei, we performed snRNA-seq on post-mortem hypothalamic tissue from CTL and early-stage AD donors, complemented by spatial transcriptomics data of the human hypothalamus. This approach allowed us to dissect cell-type-specific contributions to hypothalamic pathology, revealing key molecular features related to inflammatory response and circadian rhythms distinguishing between cells from CTL and AD donors by machine learning. Further analysis with trajectory inference and differential expression identified striking vulnerabilities in microglia, GABAergic/inhibitory neurons, and specifically circadian rhythms and sleep-regulating neurons predicted to be from SCN and DMH. Consistent with other brain regions, microglia, which are widely recognized mediators of brain aging and neurodegeneration^13,50,51^, emerged as particularly prone to transcriptional reprogramming in the hypothalamus in early AD. They displayed enriched signatures for proliferation (e.g., G2M checkpoint regulation) and metabolic remodeling (e.g., mTORC1 signaling, cholesterol homeostasis). Using trajectory inference^52,53^, we identified a continuum of microglial states transitioning from aging-related profiles to AD-associated profiles, with homeostatic microglia progressively shifting toward DAM phenotypes. These observations reinforce the concept that microglia may pivot from supportive roles toward more inflammatory or dysregulated phagocytic states in early AD, potentially exacerbating tissue damage. In parallel, we detected increased senescence and NFT pathway signatures in microglia, leukocytes, and endothelial cells^46,47^. This further suggests that diverse cell populations contribute to a proinflammatory and proteostasis-stress environment in the AD hypothalamus, while neurons primarily accumulate NFTs in later disease stages. Additionally, the molecular analysis implies the biological underpinnings of the volumetric changes captured by the imaging studies, such as the respective roles of microglial inflammation, mTORC1 signaling in oligodendrocytes, and intracellular NFTs.

In neurons, our single-cell data pinpointed a pronounced dysregulation in hypothalamic GABAergic cells, particularly those implicated in neuropeptide signaling and GPCR pathways, raising the possibility that crucial metabolic and circadian circuits become perturbed early in the disease process. Of particular note is the SCN, the brain’s master circadian clock^60^, where vasopressin signaling pathways were altered in our human AD samples. Moreover, the SCN and DM were also identified as being particularly affected in our volumetric analysis by aging and cognitive decline. Given that circadian rhythms disruption is a well-established, often prodromal, feature of AD^16,61^, these results suggest a mechanistic link whereby age-related SCN dysfunction and associated neuropeptide imbalances could drive both sleep disturbances and metabolic dysregulation, thereby contributing to the development and progression of AD.

Despite these important findings, some limitations warrant consideration. First, our volumetric analysis involves a relatively small number of individuals with biomarker-positive AD, which might limit its sensitivity to detecting AD-related effects. Second, our human donor cohort was relatively small, reflecting the inherent challenges in obtaining well-preserved hypothalamic tissues with thorough clinical and neuropathological annotations. Although we implemented strategies to minimize confounders (e.g., controlling for post-mortem interval, RNA integrity, and sex in specific analyses), larger datasets are needed to fully confirm the specificity and temporal progression of the identified subregion-and cell-type-level changes^62^. Third, due to the high cost of single-cell sequencing, cells captured in HypoAD were a subset of the hypothalamic subregions manually enriched for, leaving some rare cell types that change with AD undiscovered. Forth, our single-cell transcriptomic approach provides a snapshot of gene expression; whether these transcriptional signatures translate directly into the proposed functional consequences, such as disrupted circadian rhythms or altered metabolic rates, remains to be explicitly evaluated through complementary electrophysiological, behavioral, and in vivo imaging assays.

In summary, our integrative volumetric and single-cell atlas of the human hypothalamus provides a high-resolution framework for elucidating how aging and AD pathology differentially affect specific hypothalamic subregions and cell types. We highlight notable vulnerabilities in GABAergic/inhibitory neurons, SCN and DMH circadian regulators, and inflammatory microglial cascades, underscoring their potential roles in instigating early hypothalamic dysfunction. These findings converge on the central themes of neuroinflammation and proteostasis stress, suggesting that subregion-specific susceptibility, driven by distinct cellular responses, may be a key factor in disease progression. Moving forward, the identified pathways and circuits present valuable targets for therapeutic strategies aimed at preserving neuropeptide-GPCR signaling, maintaining circadian homeostasis, and modulating microglial activation, thereby offering potential avenues to mitigate hypothalamic dysfunction in Alzheimer’s disease.

## Methods

### Volumetric analysis of imaging data

De-identified high-resolution MRI scans were obtained from the study by Goyal et al. (2023)^63^. The native brain images were subsequently preprocessed and registered to a high-resolution template previously generated and described by Neudorfer and Germann et al. (2020)^25^. Given that the dataset did not have predefined group structures for all desired comparisons, we employed a GLM to assess the associations between hypothalamic subregion volumes (summed across left and right hemispheres) and various demographic and clinical variables. Specifically, we modeled the natural logarithm of each subregion’s volume (log(subregion volume)) as a function of scaled age at MRI, sex, cognitive status, amyloid-β deposition status, and other relevant covariates:

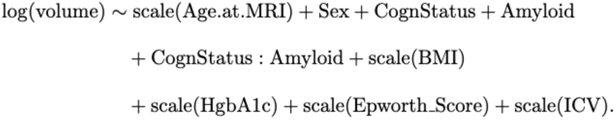

P-values for individual variables from the GLM analyses were corrected for multiple comparisons using the Benjamini–Hochberg (BH) procedure to control the false discovery rate.

To specifically assess the effect of AD on subregion volumes, participants were categorized based on their cognitive status with Clinical Dementia Rating (CDR) score (e.g., normal with CDR 0 or impaired with CDR>0) and amyloid-β PET or CSF biomarker status (negative or positive for amyloid-β burden). Individuals meeting criteria for both cognitive impairment and positive amyloid-β status were designated as the AD group. Conversely, participants without cognitive deficits and with negative amyloid-β status were classified as the non-AD (control) group. To minimize potential confounding from intermediate or mixed pathology profiles, only individuals clearly falling into these AD or non-AD categories were included in these specific comparative analyses.

To isolate the volumetric changes associated with AD from potential confounding by age and sex, we implemented a propensity score matching procedure using the MatchIt R package (matchit(AD ∼ Sex + Age.at.MRI, method = “nearest”)). First, a logistic regression model estimated each participant’s propensity score for being in the AD group based on their sex and age at MRI. A nearest-neighbor algorithm then used these scores to generate matched pairs of AD and non-AD individuals with balanced distributions of these key covariates. This approach enabled direct volumetric comparisons between the groups using paired Wilcoxon signed-rank tests with BH correction.

### Post-mortem human brain tissues

Unfixed, flash-frozen half-hypothalamus tissues were obtained from the Netherlands Brain Bank (https://www.brainbank.nl), a registry that obtains written informed consent for the use of material and clinical data for research purposes. This consent procedure was approved by the Ethical Committee of the VU University Medical Center (Vrije Universiteit, Amsterdam, The Netherlands), independently and prior to this study. Tissue blocks were stored at -80 °C and subsequently processed in a cryostat to enrich for the hypothalamic subregions of interest.

### Statistics and reproducibility

No statistical methods were used to predetermine sample sizes but our sample sizes are similar to those reported in a previous publication using single-cell RNA-seq on brain tissue^64^. Individual nuclei were excluded from analysis based on quality control metrics (feature count and mitochondrial read count). Data collection and analysis were not performed blind to the conditions of the experiments.

### snRNA-seq

Nuclei extraction was performed using the Nuclei PURE Prep kit (Millipore Sigma) according to the manufacturer’s instructions. All the procedures were performed in the cold room before the 45 min spin down. Tissue was transferred using a transfer pipette into a refrigerated Dounce homogenizer with 5 ml lysis solution following kit instructions. Tissue was homogenized with the Dounce B and the lysate was transferred into a 15 ml Falcon tube through a 40 μm filter. The sucrose purification step was performed with the modifications that half the volume of all reagents was used to account for the small tissue sample sizes, an SW34 rotor was used and samples were spun for 45 min at 30,000g (13,000 r.p.m.) at 4 °C.

Nuclei were counted using a hemocytometer and 10,000 cells per sample were loaded onto the Chromium Single Cell 3’ Chip (10X Genomics) and processed with the Chromium Controller (10X Genomics). Samples were sequenced at GENEWIZ on an Illumina NovaSeq, with a target of 50,000 reads per sample.

### Quality control, data processing and analysis

Sequence alignment for human snRNA-seq data (including the dataset from Siletti et al.) was performed against the GRCh38 (2020) reference genome. These alignments were executed using CellRanger (10x Genomics; v7.1.0 for human) with the --include-introns flag, which is essential for capturing nuclear RNA.

Following alignment, systematic background noise, such as ambient RNA, was removed from the count matrices using CellBender (v0.3.0)^65^. For quality control (QC), outlier cells were identified based on established criteria. Specifically, cells were flagged as outliers if they exceeded 5 median absolute deviations (MADs) for any of the following QC metrics: log1p_total_counts, log1p_n_genes_by_counts, or pct_counts_in_top_20_genes. Additionally, cells with a mitochondrial transcript percentage of ≥ 10 were excluded.

Potential doublets (two or more cells captured as a single droplet) were then identified and filtered out using scDblFinder^66^. After these QC steps, data from different individuals were integrated using scVI (single-cell Variational Inference)^32^ to correct for batch effects. Subsequent data visualization and downstream analyses were performed using tools from the Seurat^67^ and Scanpy^68^ ecosystems.

DE analysis was performed using a pseudobulk approach with the LRTin edgeR^69^. Genes were significant if the BH-adjusted p-value was below 0.05.

GSEAwas performed using the fgsea package^70^ using the hallmark gene set list and KEGG and Reactome gene set list from MSigDB^70,71^. For GSEA input, genes within each cluster were ranked by their log_2_FC values, determined from MAST analysis^72^. The GSEA was then performed using the fgseaMultilevel() command with default settings. Gene sets were considered significantly enriched if the BH-adjusted p-value was < 0.1.

### Embedding analysis with UCE

To contextualize our data within a broader reference, we utilized a Universal Cell Embedding (UCE) foundation model. UCE projects single-cell gene expression profiles into a universal embedding space that is agnostic to technical covariates like batch effects or protocol differences^73^. We implemented the 33-layer version of UCE as specified in its official repository (https://github.com/snap-stanford/UCE), using Python (v3.9.0) and PyTorch (v2.1.1), using the publicly available pretrained model weights.

We then used UCE to project our CTL and AD snRNA-seq datasets into a shared embedding space alongside a publicly available, community-curated human brain atlas containing over 25 million cells^73^. The resulting embeddings were visualized in a 2D UMAP space to benchmark our newly generated data against this large-scale public reference.

### Machine Learning Model Training and Feature Ranking

To identify predictive features for the classification task distinguishing cells from AD versus control individuals, we developed and applied a machine learning pipeline using Python (v3.10.9) and the Scikit-learn library (v1.6.1). The pipeline involves four main stages: subsetting the data, performing a nested cross-validation, and finally, extracting and ranking feature weights for GSEA.

Data Subsetting. For cell-type-specific models, the full dataset was filtered to include only cells corresponding to a single target cell type (e.g., neuron, astrocyte), based on pre-defined annotations.

Nested Cross-Validation. We employed a nested cross-validation (CV) strategy to ensure robust hyperparameter tuning and unbiased performance evaluation. The data for all cells, as well as for each of the top five most abundant cell types, was split into seven outer folds based on the sample batch of origin (leave-one-batch-out). For each outer fold, the remaining six batches were used for an inner leave-one-batch-out CV to select the optimal hyperparameters for an elastic net logistic regression model. The 5×5 hyperparameter grid included a logarithmic search for the regularization parameter C (from 10⁻² to 10²) and a linear search for the L1 ratio (from 0 to 1). The model with the best average inner-fold area under the receiver operating characteristic curve (AUROC) was then retrained on all six training batches and evaluated on the held-out test batch.

Feature Weight Extraction and Ranking. Following the outer CV, we extracted feature weights (coefficients) from the models on each of the seven folds. We used these weights to generate two distinct feature ranking lists for each cell type. The first, a “raw-weight” rank, ordered features by their signed coefficients to indicate a positive or negative association with the prediction. The second, an “absolute-weight” rank, used the absolute value of the coefficients to measure overall predictive importance, regardless of direction. To establish a final, robust ranking for each, we averaged each list across the seven folds.

To identify features that were consistently important across all cell types, we first calculated a “shared score” for each gene. This score was defined as the mean of the gene’s absolute-weight rank across the five cell types. The directionality of this association (positive or negative) was based on the sign of the gene’s coefficient in the raw-weight rank.

To quantify cell-type specificity, we then calculated a specificity score using the formula:

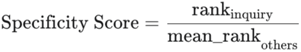

where rank_inquiry_ is the feature’s rank in the cell type of interest and mean_rank_others_ is its average rank across the remaining cell types.

GSEA. To identify biological pathways associated with our model’s predictions, we performed pre-ranked GSEA using the gseapy (v1.1.9) Python package. For each cell type, the full list of features was ranked using the mean raw weight. This ranked list was tested for enrichment against the KEGG pathway database (v2021), Hallmark (v2020), and GO Biological Process (2021) from the Molecular Signatures Database (MSigDB). Significance was determined using a permutation-based test (1,000 permutations), and pathways with a BH-adjusted p-value< 0.05 were considered significantly enriched.

### Trajectory inference and analysis using Monocle3

To infer the aging process for the microglia cluster (n = 51,420 nuclei) generated in Seurat, we applied Monocle3^51,52^. Monocle3 employs dimensionality reduction to position cells in a lower-dimensional space, can correct for batch effects using mutual nearest neighbor alignment, and subsequently connects cells to construct trajectories in a semi-supervised manner.

For our analysis of the microglia/macrophage cluster, we directly used the integrated Seurat object, bypassing any additional batch correction or dimensionality reduction steps within Monocle3 itself, as these had been addressed in the prior Seurat processing. After subsetting the microglia cluster, the root of the trajectory was defined by selecting the principal graph node most enriched for cells originating from young individuals. Spatial differential expression analysis along this inferred trajectory was then performed using Moran’s *I* test, as implemented in Monocle3. Genes with a BH-adjusted p-value < 0.01 were considered significantly trajectory-dependent, resulting in the identification of 13,104 such genes.

### Functional enrichment analysis

Functional enrichment analysis of the 13,104 t-DEGs, grouped into individual expression groups identified through co-expression analysis, was performed using Enrichr (v3.0)^74^. This analysis utilized the “GO_Biological_Process_2018” gene set database. Statistically significant enriched Gene Ontology (GO) terms were identified based on a BH-adjusted p-value < 0.01. For visualization purposes, the top ten significantly enriched terms per group were selected in a dot plot.

To decode and visualize the dynamic expression patterns of the t-DEGs (n=13,104) along the inferred pseudotime trajectory, we employed the RVAgene package (v1.0)^57^ in Python (v3.9.6) with PyTorch (v1.9.0). Prior to input into RVAgene, expression values for each gene were averaged within discrete pseudotime bins and subsequently rescaled to a range of [-1, 1]. The RVAgene neural network was configured with two hidden layers (48 nodes per layer) and two latent variable dimensions. The smoothed, reconstructed expression trajectories of the t-DEGs generated by RVAgene were then used to create the summary heatmap.

### Spatial transcriptomics analysis with Cell2location

Cell2location is a Bayesian model that uses snRNA-seq cell-type signatures to infer cell types in Visium spatial transcriptomics by decomposing mRNA counts in each Visium voxel into cell types^59^. We performed the three main steps in the cell2location workflow: estimate reference expression signatures of cell types using our dataset, map the learned cell type signatures onto the slides and perform downstream analysis. We used the default parameters to train the cell2location model.

## Acknowledgements

Funding for the acquisitions and managing of the imaging dataset in Washington University in Saint Louis was provided by NIH/NIA R01AG053503, R01AG057536, and RF1AG073210. We are particularly grateful to research participants for their altruism. We also acknowledge the Neuroimaging Laboratory, Knight Alzheimer’s Disease Research Center (particularly Drs. JC Morris and TLS Benzinger), and the imaging staff for making the dataset collection possible. Some of the MRI sequences used in the AMBR dataset were obtained from the Massachusetts General Hospital.

This work was supported by a Carney Institute Innovation Award to A.E.W.. D.Y. was supported by the NIH/NIA F99/K00 award AG083292. K.R.B. was supported by the Blavatnik Family Graduate Fellowship in Biology and Medicine. K.H.H. was funded by Neustein Graduate Fellowship from the Carney Institute for Brain Science at Brown University. We are grateful to the donors of postmortem tissue to the Netherlands Brain Bank. We thank Dr. Christoph Schorl (Brown Genomics Core) for support with sample and library quality control, and members of the Webb laboratory and Noble laboratory for providing critical feedback. This research was conducted using computational resources and services at the Center for Computation and Visualization at Brown University. Schematic figures were made with BioRender.

## Author contributions

D.Y. and A.E.W. conceptualized, designed, and coordinated the study. D.Y. conducted all experiments unless otherwise specified. J.G. performed hypothalamic subregion segmentation. G.M. and R.S. performed the embedding analysis. S.K.D. performed senescence analysis supervised by M.E.O. and H.Z. I.V. provided guidance on tissue dissection and sample selection. K.R.B. and K.H.H. assisted with tissue processing. A.C.J. contributed to cell type annotation. W.S.N. provided guidance on machine learning and interpretation. M.S.G. and A.G.V. provided the MRI dataset and guided the volumetric analysis. D.Y. and A.E.W. wrote the manuscript, with all authors providing critical feedback.

## Data availability

The raw data for snRNA-seq of young human hypothalamus were accessed from https://data.nemoarchive.org/biccn/grant/u01_lein/linnarsson/transcriptome/sncell/ <u>10x_v3/human/raw/</u>. The spatial transcriptomics data of the human hypothalamus were accessed from https://www.ebi.ac.uk/biostudies/arrayexpress/studies/E-MTAB-11114.

## Competing Interests

The authors declare no competing interests.

---

**Extended Data Figure 1.**
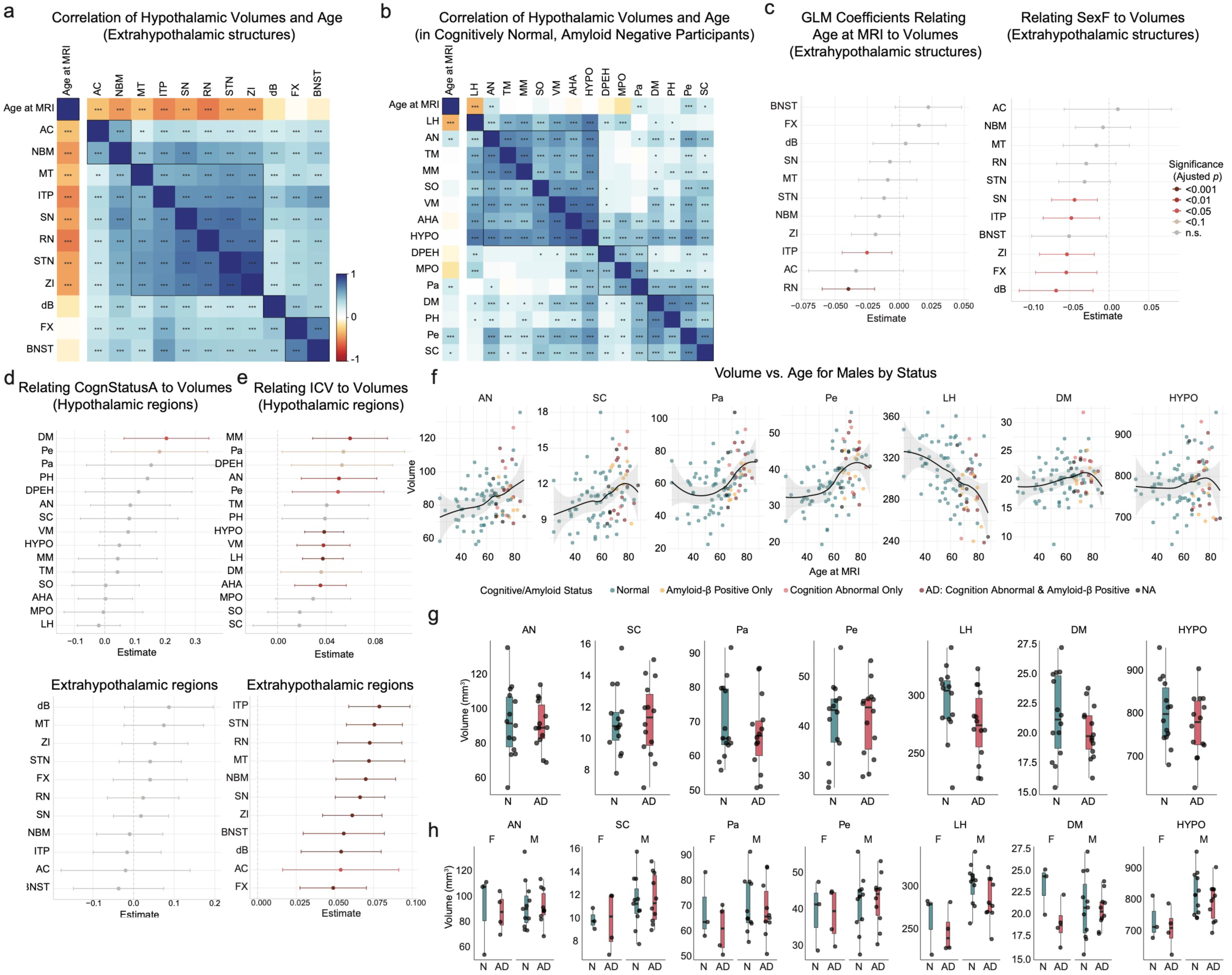
Analysis of hypothalamic and extrahypothalamic structure volumes. **a,** Pearson correlation heatmap of extrahypothalamic structure volumes with age at MRI. Blue and red indicate positive and negative correlations, respectively. Significance was determined after BH correction (\**p* < 0.05, \*\**p* < 0.01, \*\*\**p* < 0.001). **b,** Same as in (**a**), but for amyloid-β-negative and cognitively normal individuals. **c,** Forest plots showing GLM coefficients for the effects of age at MRI (left) and biological sex being female (right) on extrahypothalamic structure volumes. Points represent coefficient estimates and error bars represent 95% confidence intervals. Significance is based on BH-corrected p-values. **d,** Same as in (**c**), relating cognitive status being abnormal to volumes of hypothalamic (top) and extrahypothalamic (bottom) subregions. **e,** Same as in (**c**), relating ICV to volumes of hypothalamic (top) and extrahypothalamic (bottom) subregions. **f,** Scatterplots depicting age-related volumetric changes (mm^3^) in selected hypothalamic subregions and the total hypothalamus in males. Each point represents an individual and is colored by their combined cognitive and amyloid-β status. A locally estimated smoothing (LOESS) curve (black line) and its 95% confidence interval (shaded area) are shown for each plot. **g-h**, Boxplots comparing volumes of hypothalamic subregions and the total hypothalamus between males and females within cognitively normal (N) and AD groups. Data were propensity-score matched for age using nearest-neighbor matching. Boxplots display the median, interquartile range (IQR), and whiskers extending to 1.5 times the IQR. **(g)** Data are shown for males and females combined. **(h)** Data are shown with males and females separated.

**Extended Data Figure 2.**
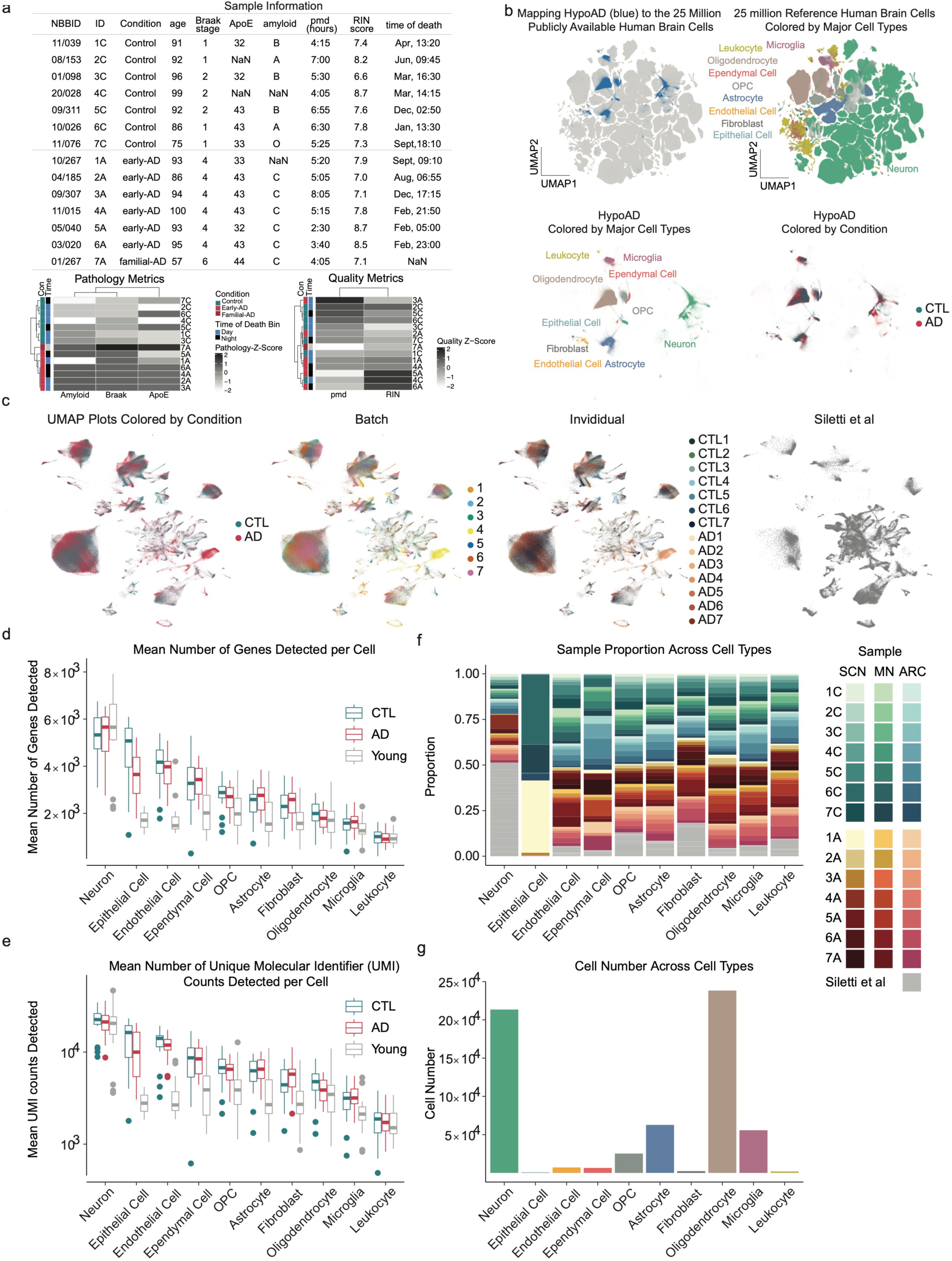
Quality control and sample information. **a**, Sample metadata. A table summarizes the condition, age, Braak stage, *APOE* status, amyloid-β status, postmortem delay (PMD), RNA integrity number (RIN), time of death, and presence of documented sleep changes for each sample. Accompanying heatmaps show samples clustered by pathology metrics (left) and quality metrics (right), with annotation bars indicating condition, time of death, and sleep change status. **b**, UMAP showing nuclei from this study embedded within a reference atlas of 25 million publicly available human brain cells, colored by major cell types and condition. **c**, UMAP of the integrated dataset following scVI batch correction, with nuclei colored by condition, batch, individual, and the Siletti et al dataset. **d**, Boxplots comparing the number of genes detected per nucleus in neurons, glial cells, and immune cells across young control, aged control (CTL), and AD samples. **e**, Boxplots showing the distribution of unique molecular identifier (UMI) counts per nucleus across major cell types and conditions. **f**, Stacked bar plots showing the cell type composition for samples from different hypothalamic subregions (SCN, MN, ARC). **g**, Bar plot quantifying the total number of nuclei for each cell type in the final dataset.

**Extended Data Figure 3.**
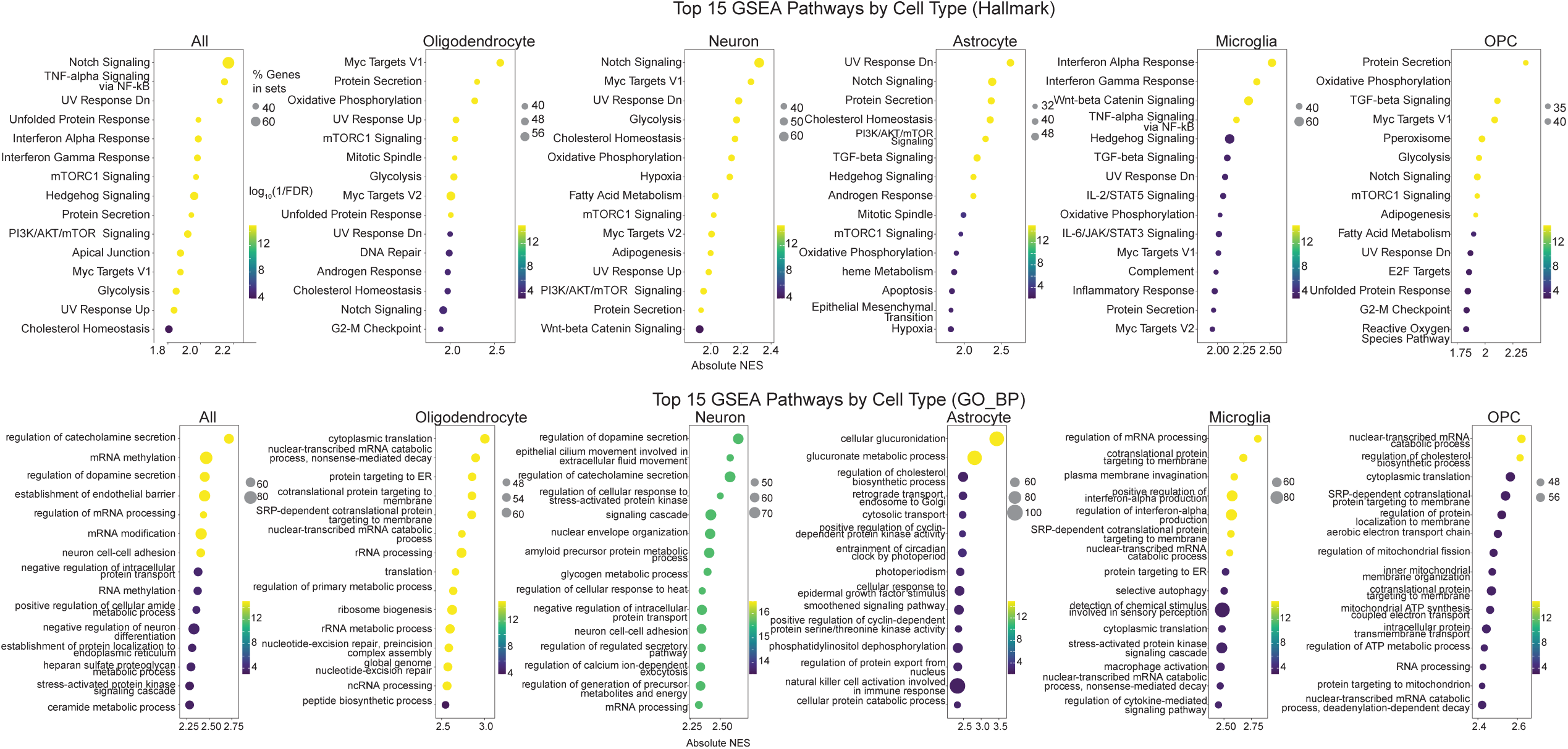
GSEA reveals biological pathways associated with predictive AD features. Selected significantly enriched Hallmark (left) and gene ontology (GO) biological processes (BP) (right) pathways from GSEA. The analysis was performed on genes ranked by their signed model weights. The dot plot shows the top pathways for the all-cell model and for each of the five cell-type-specific models. The x-axis represents the absolute normalized enrichment score (NES), dot size represents the number of genes in a pathway, and color indicates to the significance (−log10(FDR)).

**Extended Data Figure 4.**
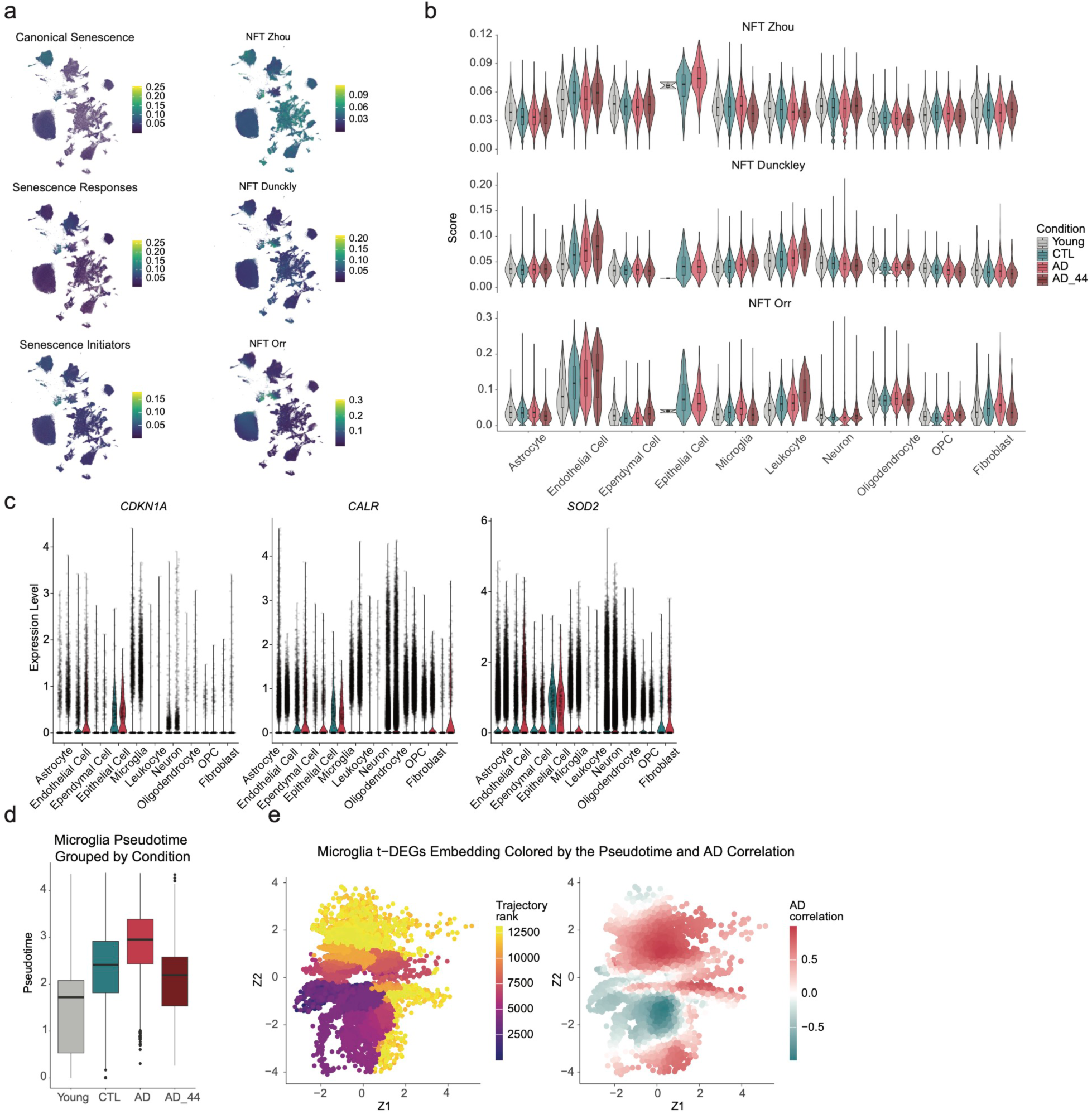
Transcriptional changes in HypoAD. **a**, UMAP projections illustrating the expression scores for three senescence-related and three neurofibrillary tangle-related gene signatures across all nuclei. **b**, Violin plots comparing the expression scores of three neurofibrillary tangle-related signatures across major cell types. **c**, Violin plots showing the expression of key senescence-related genes (*CDKN1A* – canonical, *CALR* – Responses, and *SOD2* – Initiators) across major cell types. **d**, Boxplots comparing the microglial pseudotime distribution across young, control (CTL), AD, and a familial AD (AD_44) case. **e**, UMAP embeddings of microglia. Left: Nuclei are colored by their rank along the pseudotime trajectory. Right: The same nuclei are colored by the correlation of their expression profile with AD status.

**Extended Data Figure 5.**
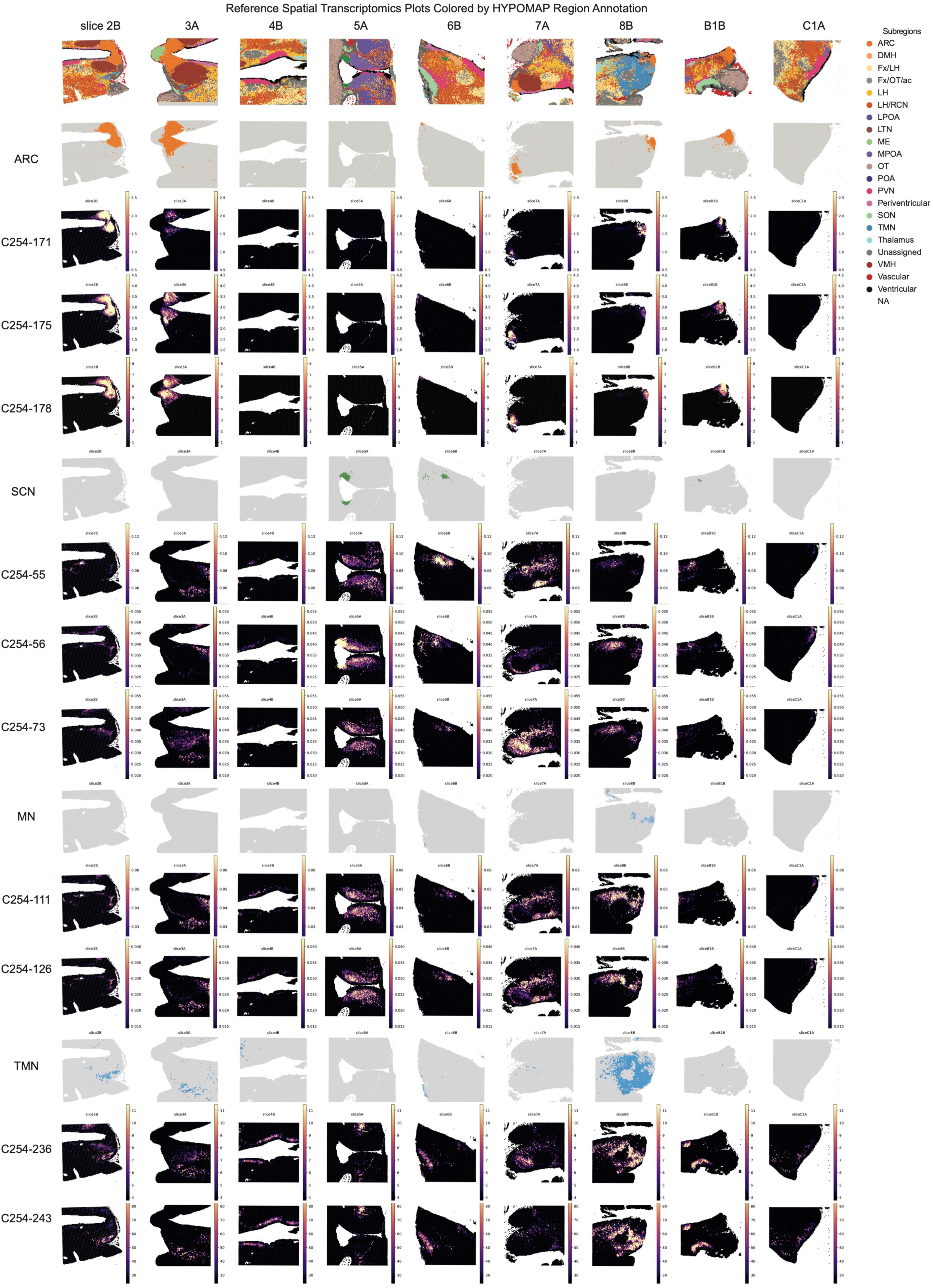
Spatial mapping of neuronal subtypes using Visium data.

**Extended Data Figure 6.**
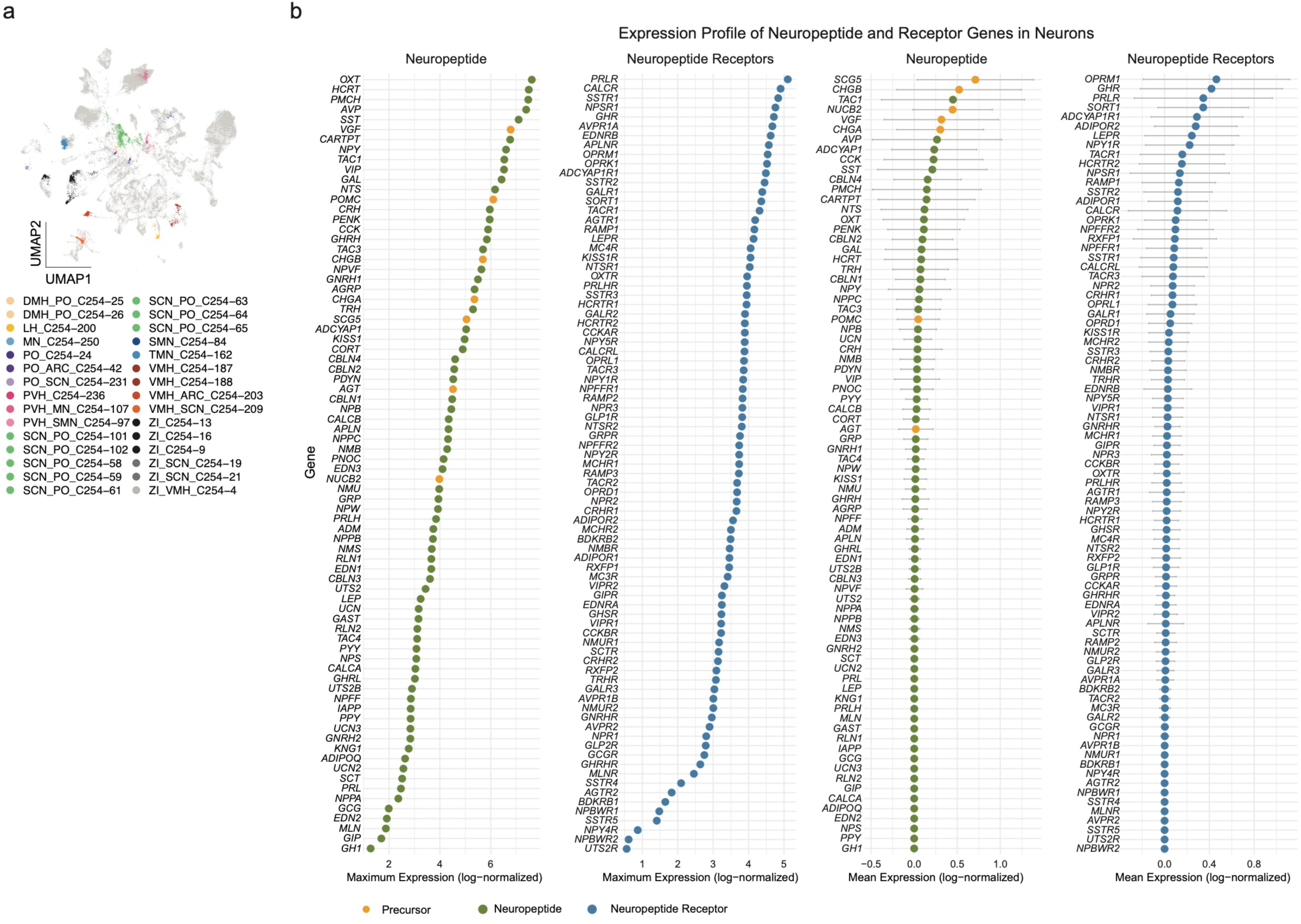
Analysis of neuronal subclusters with the most DEGs and neuropeptide expression. **a.** UMAP of the top 30 neuronal subclusters with the highest number of DEGs. **b.** Dot plots showing maximum and mean expression levels for neuropeptide genes and their corresponding receptors in neurons. For mean expression plots, error bars represent the standard deviation across all neurons.

**Supplementary Table 1:** Abbreviations for hypothalamic and extrahypothalamic subregions in MRI and snRNA-seq analyses.

**Supplementary Table 2:** Pearson correlation matrix of subregion volumes with age, including BH-corrected p values.

**Supplementary Table 3:** Results of GLMs for individual subregions.

**Supplementary Table 4:** GSEA results of predictive features from the machine learning models.

**Supplementary Table 5:** Results of edgeR-LRT differential expression analysis for major cell types.

**Supplementary Table 6:** Results of Moran’s *I* test for identifying trajectory-varying genes in microglia.

**Supplementary Table 7:** Results of edgeR-LRT differential expression analysis for neuronal subtypes.

## Notes

### Competing Interest Statement

The authors have declared no competing interest.

